# Random Walk With Restart on Multiplex and Heterogeneous Biological Networks

**DOI:** 10.1101/134734

**Authors:** Alberto Valdeolivas, Laurent Tichit, Claire Navarro, Sophie Perrin, Gaëlle Odelin, Nicolas Levy, Pierre Cau, Elisabeth Remy, Anaïs Baudot

## Abstract

Recent years have witnessed an exponential growth in the number of identified interactions between biological molecules. These interactions are usually represented as large and complex networks, calling for the development of appropriated tools to exploit the functional information they contain. Random walk with restart is the state-of-the-art guilt-by-association approach. It explores the network vicinity of gene/protein seeds to study their functions, based on the premise that nodes related to similar functions tend to lie close to each others in the networks.

In the present study, we extended the random walk with restart algorithm to multiplex and heterogeneous networks. The walk can now explore different layers of physical and functional interactions between genes and proteins, such as protein-protein interactions and co-expression associations. In addition, the walk can also jump to a network containing different sets of edges and nodes, such as phenotype similarities between diseases.

We devised a leave-one-out cross-validation strategy to evaluate the algorithms abilities to predict disease-associated genes. We demonstrate the increased performances of the multiplex-heterogeneous random walk with restart as compared to several random walks on monoplex or heterogeneous networks. Overall, our framework is able to leverage the different interaction sources to outperform current approaches.

Finally, we applied the algorithm to predict genes candidate for being involved in the Wiedemann-Rautenstrauch syndrome, and to explore the network vicinity of the SHORT syndrome.

The source code and the software are freely available at: https://github.com/alberto-valdeolivas/RWR-MH.

## 1 INTRODUCTION

Recent years have witnessed the accumulation of physical and functional interactions between biological macromolecules. For instance, protein-protein interactions (PPI) are nowadays screened at the proteome scale for many organisms, including humans, revealing thousands of physical interactions between proteins. Interaction data are commonly represented as networks, in which the nodes correspond to genes or proteins, and the edges to their interactions. The availability of large-scale PPI networks led to the application of graph theory-based approaches for their exploration, with the ultimate goal of extracting the knowledge they contain about cellular functioning. These methods exploit the tendency of functionally-related proteins to lie in the same network neighborhood. For instance, clustering algorithms allow identifying communities of proteins participating in the same biological processes (Brohée and van Helden, 2006; Katsogiannou et al., 2014; Chapple et al., 2015; Arroyo et al., 2015) and guilt-by-association strategies explore topological relationships to predict protein cellular functions (Schwikowski et al., 2000).

Network-based guilt-by-association strategies, in particular, have been widely used to identify new disease-associated genes. The first approaches were parsing the direct interactors of disease proteins in a PPI network (Oti et al., 2006). Then, more elaborated algorithms were developed, computing the shortest path distances between candidates and known disease proteins (Franke et al., 2006; George et al., 2006). But algorithms able to exploit the global topology of networks, such as network propagation or random walk algorithms, were finally shown to largely outperform initial methods to identify disease genes (Vanunu et al., 2010; Köhler et al., 2008).

Random walks were indeed first developed to explore the global topology of networks. They simulate a particle that iteratively moves from a node to a randomly selected neighboring node (Lovász, 1993). The idea of restart, which led to the Random Walk with Restart (RWR) algorithm, was first introduced for internet search engines. The objective was to rank the relevance of web pages by reproducing the behavior of a simulated Internet user. The user surfs randomly from a web page to another thanks to the hyper-links, but he can also restart the navigation in a new arbitrary web page. Thereby, depending on the topological structure of the pages and hyper-links, some pages will be visited more frequently than others. The number of visits is considered as a proxy measure of each web page relevance (Brin and Page, 1998). Moreover, if one forces the particle to always restart in the same node or set of nodes – called *seed(s)* -, RWR can be used to measure a proximity between the seed(s) and all the other nodes in the network (Pan et al., 2004).

RWR became the state-of-the-art guilt-by-association algorithm in network computational biology. It was initially applied, as commented above, to prioritize candidate disease genes. All the network nodes are ranked by the RWR algorithm according to their proximity to known disease-associated nodes taken as seeds (Köhler et al., 2008). Several extensions of the RWR algorithm further improved the prediction of candidate disease-associated genes, mainly by con-sidering also phenotype data (Li and Patra, 2010; Li and Li, 2012; Xie et al., 2015; Zhao et al., 2015). For instance, Li and Patra (2010) described a RWR on a heterogeneous network, built by connecting a PPI network with a disease-disease network using known gene-disease associations.

However, a common feature and limitation of these approaches is that they perform the walks in a single network of relationships between genes and proteins. Doing so, these approaches ignore a rich variety of information on physical and functional relationships between biological macromolecules. Indeed, not only PPI are nowadays described on a large-scale: immuno-precipitation experiments followed by mass-spectrometry can inform on the *in vivo* molecular complexes (Ruepp et al., 2009), pathways interaction data are cured and stored in dedicated databases such as Reactome (Fabregat et al., 2016) and Kegg (Kanehisa et al., 2008). In addition, other functional interactions can be derived, for instance from transcriptomics expression data by constructing a co-expression network, or from gene ontology annotations (Ashburner et al., 2000) by constructing a co-annotation network.

Each interaction source has its own meaning, relevance and bias: some networks contain links of high relevance (e.g., curated signaling pathways), while others contain thousands or even millions of interactions prone to noise (e.g., co-expression networks) (Didier et al., 2015). The combination of the different sources is expected to provide a complementary view on genes and protein cellular functioning (Menche et al., 2015). But networks can be combined in different ways. Generally, the different networks are merged into an aggregated monoplex network. For instance, Li and Li (2012) adapted the RWR algorithm to a network in which PPI and co-annotation interactions were aggregated. However, aggregating interactions networks sources as a single networks dismisses the individual networks topologies and features. In this context, the multiplex framework offers an interesting alternative. Collections of networks sharing the same nodes, but in which the edges belong to different categories or represent interactions of a specific nature are called multiplex (alt. multi-slice, multi-layer) networks (Battiston et al., 2014). In a biological multiplex network, each layer contains a different category of physical and functional interactions between genes or proteins, with its own topology.

We present here two extensions of the RWR algorithm to explore multiplex networks (RWR-M) and multiplex-heterogeneous networks (RWR-MH). We constructed a multiplex network composed of three layers of physical and functional interactions between genes and proteins, and a disease-disease network based on phenotype similarities. We applied a leave-one-out cross-validation (LOOCV) strategy to compare the RWR-M and RWR-MH algorithms to alternatives, including RWR on monoplex networks, aggregated networks and heterogeneous-only networks. We showed that considering many interaction sources through a multiplex-heterogeneous network framework enhances remarkably the performances of disease-gene prioritization. Finally, we applied the RWR-MH algorithm to predict candidate genes for being implicated in the Wiedemann-Rautenstrauch syndrome (WRS), whose responsible gene(s) remain unknown. We also explored the network vicinity of the SHORT syndrome (SS) and its associated gene, *PIK3R1*, and unveiled associated syndromes and pathways.

## 2 MATERIALS AND METHODS

### 2.1 Random walks on graphs

Let us consider an undirected graph, *G* = (*V, E*). An imaginary particle starts a random walk at an initial node *v*_0_ ∈ *V*. Considering the time is discrete, *t* ∈ N, at the *t*-th step the particle is at node *v*_*t*_. Then, it walks from *v*_*t*_ to *v*_*t*+1_, a randomly selected neighbor of *v*_*t*_, in the graph *G* (Lovász, 1993). Therefore, we can write: ∀*x, y* ∈ *V*, ∀*t* ∈ ℕ

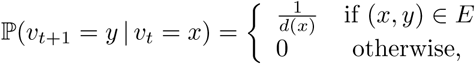

where *d*(*x*) is the degree of *x* in the graph *G*. Defining *p*_*t*_(*v*) as the probability for the random walk to be at node *v* at time *t*, we can describe the evolution of the probability distribution, **p**_*t*_ = (*p*_*t*_(*v*))_*v∈V*_, with the equation:

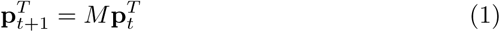

where *M* denotes a transition matrix that is the column normalization of the adjacency matrix of the graph *G*.

The stationary distribution, solution of the equation 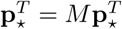, represents - if it exists - the probability for the particle to be located at a specific node for an infinite amount of time.

In order to explore the web, Brin and Page (1998) extended the classical random walk by introducing the possibility for the walk to restart. In this case, at each step, the particle can walk from its current node to either any of its neighbors or restart by jumping to any randomly selected node in the graph. This avoids the walk to be trapped in a dead end node, and assures the existence of the stationary distribution (Langville and Meyer, 2004).

Moreover, we can restrict the restart of the particle to specific set of node(s), called seed(s). At each iteration of the algorithm, the particle can restart in the seed(s) with a defined restart probability, *r* ∈ (0, 1) (Pan et al., 2004). Doing so, the particle will explore the graph focusing on the neighborhood of the seed(s). The stationary distribution is considered as a measure of the proximity between the seed(s) and all the other nodes in the graph. Formally, based on (1), RWR equation can be defined as:

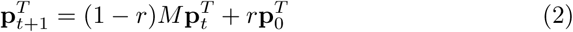

The vector **p**_0_ is the initial probability distribution. Therefore, in **p**_0_, only the seed node(s) have values different from zero. After several iterations, the difference between the vectors **p**_*t*+1_ and **p**_*t*_ becomes negligible, the stationary probability distribution is reached and the elements in these vectors represent a proximity measure from every graph node to the seed node(s). In this work, iterations are repeated until the difference between **p**_*t*_ and **p**_*t*+1_ falls below 10^−10^, as in Li and Patra (2010); Erten et al. (2011); Zhao et al. (2015).

We set here the global restart parameter to *r* = 0.7, as in previous studies (Köhler et al., 2008; Li and Patra, 2010; Li and Li, 2012; Smedley et al., 2014; Zhao et al., 2015). This value of the restart parameter is kept in all the RWR algorithms.

### 2.2 Random walks on multiplex graphs

#### 2.2.1 Definition

A multiplex graph is a collection of *L* undirected graphs, considered as layers, sharing the same set of *n* nodes (Kivelä et al., 2014). Each layer *α*, *α* = 1, …, *L*, is defined by its *n × n* adjacency matrix *A*^[*α*]^ = (*A*^[*α*]^(*i, j*))_*i,j*=1,…, *n*_, where *A*^[*α*]^(*i, j*) = 1 if node i and node j are connected on layer *α*, and 0 otherwise (Battiston et al., 2014). We do not consider auto-interactions (*A*^[*α*]^(*i, i*) = 0 ∀ *i* = 1, …, *n*), and 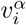 stands for the node *i* in layer *α*. A multiplex graph is characterized by its adjacency matrix:

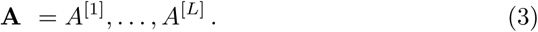

and is defined as *G*_*M*_ = (*V*_*M*_, *E*_*M*_), where:

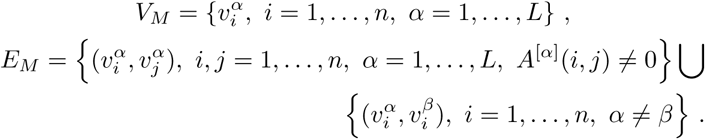

#### 2.2.2 The RWR-M algorithm: Extension of RWR to multiplex graphs

A multiplex graph contains the same set of nodes in its different layers, thereby enabling us to navigate between the layers (De Domenico et al., 2014). The particle can walk from its current node 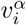 to any of its neighbors within a layer, or jump to any node 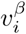 with *β* ≠ *α* (De Domenico et al., 2013), and thereby change from one to another layer, as schematically displayed in Fig. 1A.

**Figure 1:**
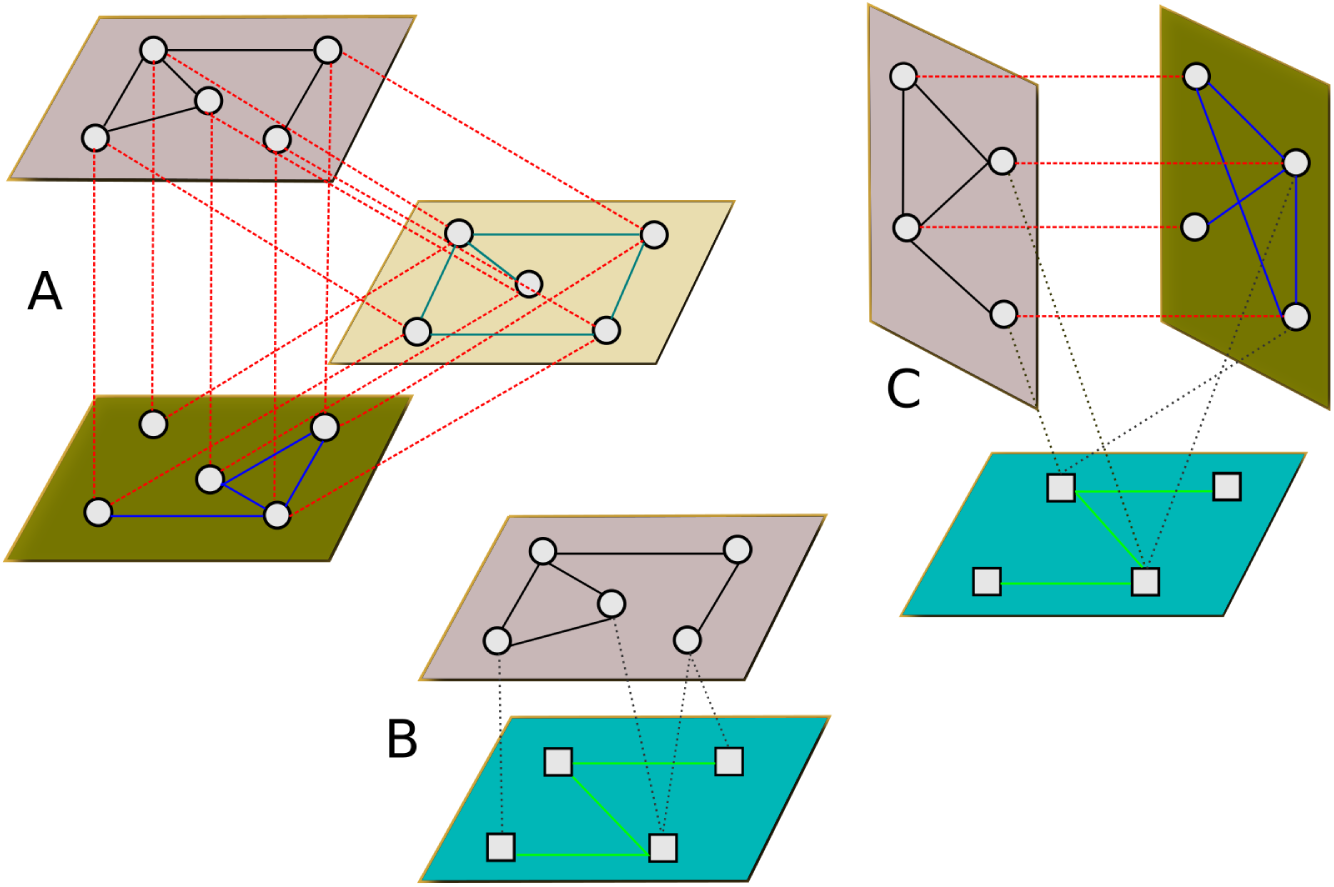
Multiplex, Heterogeneous and Multiplex-Heterogeneous graphs. **A)** A multiplex graph composed of three layers. The particle can navigate within each layer or jump to the same node in another layers. **B)** A heterogeneous graph composed of two graphs. The particle can navigate within each graph or jump to the other graph according to bipartite associations between the two different types of nodes. **C)** A multiplex-heterogeneous graph.

We can thus extend the classical RWR algorithm to multiplex graph (RWR-M) by building a *nL × nL* matrix, *A*. The matrix *A* contains the different types of transitions that the simulated particle can follow at each step, and is defined as:

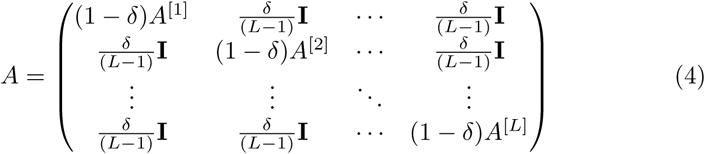

where **I** is the *n × n* identity matrix and *A*^[*α*]^ is the adjacency matrix of the layer *α*, as described in (3). The elements in the diagonal represent the potential intra-layer walks, whereas the off-diagonal elements account for the possible jumps between different layers. The parameter *δ* ∈ [0, 1] quantifies the probability of staying in a layer or jumping between the layers: if *δ* = 0 the particle will always stay in the same layer after a non-restart step. In addition, since *A*^[*α*]^(*i, i*) = 0, we avoid jumps to the same node in the same layer.

Let us denote the transition matrix *M* obtained by a column normalization of *A*. Eq. (2) in the multiplex case becomes:

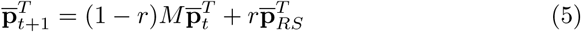

where 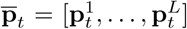 and 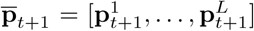, *t* ∈ ℕ, are *n × L* vectors representing the probability distribution of the particle in the multiplex graph. These vectors are composed of the probability distributions in every layer. The restart vector, 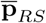, represents the initial probability distribution. We define it as 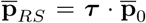, where the vector parameter ***τ*** = [*τ*_1_*, …, τ*_*L*_] measures the probability of restarting in the seed node(s) of each layer in the multiplex graph. It is to note that it is possible to tune the importance of each layer by modifying the parameter ***τ***.

As said previously, we set the global restart parameter to *r* = 0.7 for all versions of the RWR algorithm. We established an equal restart probability in all the layers, ***τ*** = (1/*L*, 1/*L*, …, 1/*L*), and we also considered an equal probability for staying in a layer or jumping between the layers, *δ* = 0.5.

The RWR-M algorithm performs iterations in Eq. (5) until the difference between 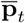 and 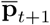 falls below 10^−10^. The stationary probability distribution is then reached, and every node is associated to *L* proximity measures, one for each layer of the multiplex graph. We computed the global score for every node as the geometrical mean of its *L* proximity measures.

For the sake of simplicity, we have considered unweighted graphs. However, the extension of the algorithms to weighted graphs is straightforward. It can be achieved by replacing the adjacency matrices (*A*^[*α*]^(*i, j*))_*i,j*=1,…,*n*_, by matrices composed of the weighted intra-layer edges (*W* ^[*α*]^(*i, j*))_*i,j*=1,…,*n*_.

### 2.3 Random walk with restart on heterogeneous graphs

#### 2.3.1 Definition

A heterogeneous graph contains two graphs with different types of nodes and edges, as well as a bipartite graph containing bipartite associations between them (Lee et al., 2013). Let us consider the graphs *G*_*V*_ = (*V, E*_*V*_) with *V* = {*v*_1_*, …, v*_*n*_}, *G*_*U*_ = (*U, E*_*U*_) with *U* = {*u*_1_*, …, u*_*m*_}, and the bipartite graph *G*_*B*_ = (*V ∪ U, E*_*B*_) with *E*_*B*_ ⊆ *V × U*. The edges of the bipartite graph only connect pairs of nodes from the different sets of nodes, *V* and *U*. We can now define a heterogeneous graph, *G*_*H*_ = (*V*_*H*_, *E*_*H*_), as:

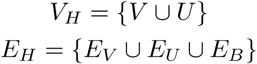

#### 2.3.2 The RWR-H algorithm: Extension of RWR to heterogeneous graphs

Li and Patra (2010) proposed a random walk with restart on a heterogeneous graph. This heterogeneous graph was composed of a PPI network, a disease-disease similarity network, and bipartite graph containing protein-disease associations. The particle walks on the PPI network, on the disease-disease similarity network, and is also allowed to jump between the two networks following the bipartite associations, as schematically displayed in Fig. 1B.

Following the approach proposed by Li and Patra (2010), let us consider the graphs defined in the previous section, *G*_*V*_, *G*_*U*_ and *G*_*B*_. We define *A*_*P* (*n×n*)_, *A*_*D*(*m×m*)_ and *B*_(*n×m*)_ as their respective adjacency matrices. These matrices are here the adjacency matrices of the PPI network, the disease-disease similarity network and the bipartite network, respectively. Therefore, the adjacency matrix of the heterogeneous network can be represented as: 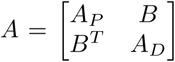, with *B*^*T*^ the transpose of the matrix *B*.

We then compute the different transition probabilities of the random walk with restart on heterogeneous graphs (RWR-H). Let 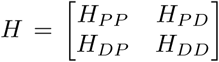 de-notes the matrix of transitions on the heterogeneous graph, where *H*_*P* *P*_ and *H*_*DD*_ describe the walks within a network, and *H*_*P* *D*_, *H*_*DP*_ describe the jumps between networks. For a given node, if a bipartite association exists, the particle can either jump between graphs or stay in the current graph with a probability given by the parameter *λ* ∈ [0, 1]. The closer *λ* is to one, the higher is the probability of jumping between networks.

Let a particle be located at the protein node *p*_*i*_ ∈ *V*. At the next step, the particle can either walk to a protein *p*_*j*_ ∈ *V* with the following transition probability:

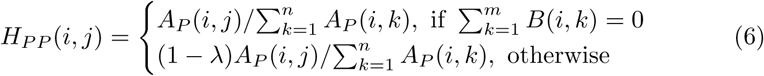

or jump through a bipartite association to the disease *d*_*b*_ ∈ *U* with a probability:

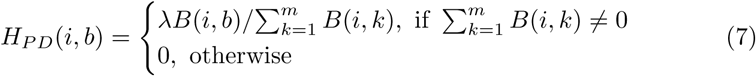

The same situation arises if a particle is located at the disease *d*_*a*_ ∈ *U*. It can walk to the disease *d*_*b*_ ∈ *U* :

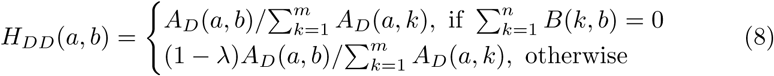

or jump to the protein *p*_*j*_ ∈ *V*:

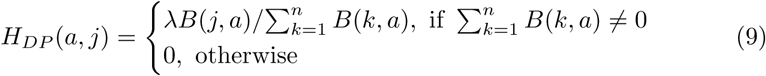

Therefore, we can write the RWR-H equation on a heterogeneous graph as:

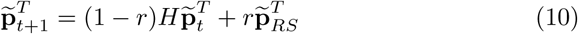

The vectors 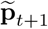,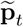 and 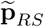 are now of dimension *n* + *m*, since the RWR-H algorithm is ranking proteins and diseases at the same time. Importantly, after a restart step, the particle can go back either to a seed protein or to a seed disease. It is to note that it is possible to tune the importance of each network by defining 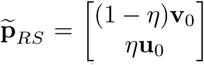, where **v**_0_ and **u**_0_ represent the initial probability distributions on the PPI and the disease-disease similarity networks given by their seed nodes. The parameter *η* ∈ [0, 1] controls the probability of restarting in each network (PPI or disease-disease). If *η <* 0.5 the particle will be more likely to restart in one of the seed proteins than in one of the seed diseases. In our work, we set both parameters *λ* and *η* to 0.5.

### 2.4 Random walk with restart on multiplex-heterogeneous graphs

#### 2.4.1 Definition

Let us consider a *L*-layers multiplex graph, *G*_*M*_ = (*V*_*M*_, *E*_*M*_), with *n × L* nodes, 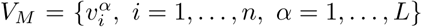. Let *G*_*U*_ = (*U, E*_*U*_) be a graph with m nodes, *U* = *{u*_1_*, …, u*_*m*_*}*. In order to build a heterogeneous graph composed of *G*_*M*_ and *G*_*U*_, we need to link the nodes in every layer of the multiplex graph *G*_*M*_ to their associated nodes in the other graph *G*_*U*_, according to their bipartite association, *E*_*B*_. Since the same nodes are present in every layer of the multiplex graph, it is necessary to have *L* identical bipartite graphs, 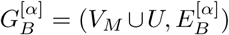 to define the multiplex-heterogeneous graph. We can then describe a multiplex-heterogeneous graph, *G*_*MH*_ = (*V*_*MH*_, *E*_*MH*_), as:

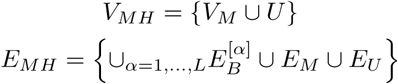

#### 2.4.2 The RWR-MH algorithm: Extension of RWR to multiplex-heterogeneous graph

We finally extended the RWR algorithm to multiplex-heterogeneous networks (RWR-MH). At a given step, let the particle be at a specific node within a layer of the multiplex graph. At the next non-restart step, the particle can either i) walk within the same layer or ii) jump to the same node in a different layer or iii) jump to the other graph if a bipartite association exists (Fig. 1C).

Let consider a multiplex graph composed of *n* gene/protein nodes and *L* layers, with an adjacency matrix *A*_*M*(*nL×nL*)_, like the one described in (4). Let also consider a disease-disease similarity graph characterized by its adjacency matrix, *A*_*D*(*m×m*)_, where *m* is the total number of diseases. The bipartite graphs with adjacency matrix 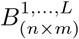 associates the gene/protein nodes in each layer of the multiplex graph to diseases. These bipartite graphs are identical for every layer of the multiplex graph, as explained previously. Therefore, we can define all of them as *B*_(*n×m*)_, and construct the bipartite adjacency matrix of the multiplex-heterogeneous graph by sticking *L* times the single bipartite graph *B*(*n×m*):

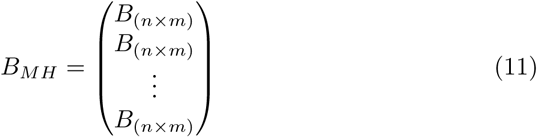

Then, we can define the global adjacency matrix of the multiplex-heterogeneous graph as 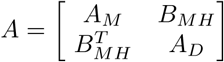, where 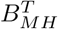 represents the transpose of *B*_*MH*_.

From this point, we can proceed in a analogous way to the one presented in section 2.3.2. We define a global transition matrix for the multiplex-heterogeneous network and calculate its components using the same equations. We just have to replace the adjacency matrix of the PPI network, *A*_*P* (*n×n*)_, by the adjacency matrix of the multiplex network *A*_*M*(*nL×nL*)_, and the bipartite adjacency matrix, *B*_(*n×m*)_, by the adjacency matrix of the bipartite graph of the multiplex-heterogeneous graph, *B*_*MH*(*nL×m*)_.

In order to apply the Eq. (10), we have to consider that the vectors 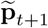, 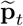 and 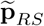 are now of dimension ((*n × L*) + *m*), since the RWR-MH algorithm is ranking *n* proteins in *L* different layers and *m* diseases at the same time. It is to note that it is possible to tune the importance of each network by defining 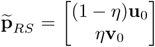, where **u** defines the initial probability distribution of the multiplex graph, as described in section 2.2.2, and **v**_0_ the initial probability distribution of the disease-disease similarity network.

### 2.5 Network sources

#### 2.5.1 Physical and Functional interactions between genes and proteins

We constructed three biological networks containing genes or proteins as nodes (genes and proteins are here considered equally): a protein-protein interaction (PPI) network, a network connecting proteins according to pathway interaction data, and a network in which the links correspond to co-expressed genes. They are obtained as described in (Didier et al., 2015), but updated from downloads on 23rd and 24th November, 2016. The PPI network contains 12 621 nodes and 66 971 edges. The Pathway network contains 10 534 nodes and 254 766 edges, and the Co-expression network is composed of 10 534 nodes connected by 1 337 347 edges (Table 1).

**Table 1:**
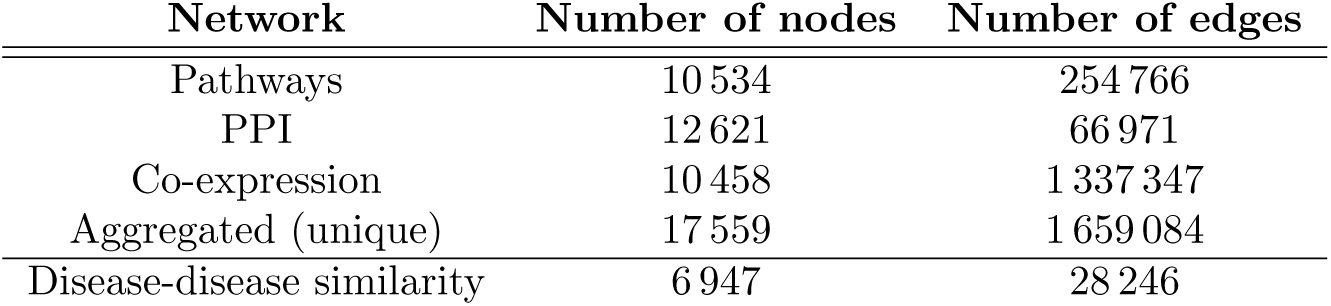
Networks used in this study, number of nodes and edges.

These networks are considered either i) independently as monoplex networks, ii) merged as an aggregated network, with nodes and edges corresponding to the union of the monoplex networks, i.e., a total of 17 559 nodes and 1 659 084 edges, or iii) as a multiplex network composed of 3 layers. In the multiplex network, the layers share the same set of nodes, also corresponding to the union of all network nodes, 17 559 nodes. The genes/proteins absent in a layer are added as isolated nodes in this layer.

#### 2.5.2 Disease-disease similarity network

We downloaded the annotation file *phenotype annotation.tab*, containing diseases and their associated phenotypes from the Human Phenotype Ontology (HPO), together with the HPO ontology graph structure (Köhler et al., 2014) on November, 2016. We kept only disease records from OMIM (Hamosh et al., 2005), and for each disease, we extracted its minimal set of HPO terms. A set of phenotypes is minimal if it describes a disease without redundancy: we considered only the deepest (i.e., the most precise) nodes in the directed ontology structure, as described by Greene et al. (2016).

The phenotype similarity between a pair of diseases can be computed by counting the number of shared phenotypes. However, some phenotypes are more relevant than others. We indeed want to consider as more similar two diseases sharing a very rare phenotype, than two diseases sharing a very common phenotype, as proposed by Westbury et al. (2015). To this goal, we estimated the relevance of each phenotype based on its frequency in the HPO database, and used the relative information content (IC), defined as follows:

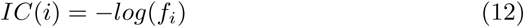

where *f*_*i*_ is the frequency of the phenotype *i* within our set of HPO diseases. The similarity between phenotypes *i* and *j* is then computed as:

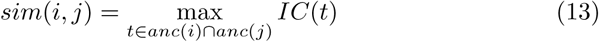

where *anc*(*i*) indicates the ancestors of the phenotype *i* in the ontology graph. Finally, the phenotype similarity between a pair of diseases *D*_*a*_ and *D*_*b*_, corresponding to two sets of HPO phenotypes, is measured by the total IC of their shared phenotypes, as described in Resnik (1999):

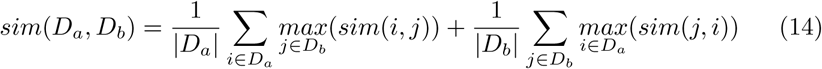

The similarity score between all pairs of diseases is computed according to Eq. (14). The disease-disease similarity network is built by linking every disease to its five nearest diseases according to this similarity score, as in Li and Patra (2010). The resulting disease-disease similarity network is composed of 6 947 diseases connected by 28 246 edges.

#### 2.5.3 Gene-Disease bipartite associations

We connected the genes in each layer of the multiplex network with the disease-disease similarity network thanks to bipartite gene-diseases associations extracted from OMIM (Hamosh et al., 2005), using BiomaRt (Durnick et al., 2012). The data were downloaded on December, 2016. The nodes in each layer of the multiplex network are connected to their related diseases, leading to L identical bipartite graphs. For each layer, we obtained 4 496 edges between genes/proteins and diseases.

### 2.6 Leave-one-out cross validation

In order to evaluate the performances of the different RWR algorithms, we designed a Leave-One-Out Cross-Validation (LOOCV) strategy. We downloaded diseases and associated genes from OMIM (Hamosh et al., 2005) (downloaded on December, 2016) and DisGeNET v4.0 (Piñero et al., 2016) (associations with a score greater than or equal to 0.15, downloaded on December, 2016). Only diseases associated to at least two genes are considered. Each gene is removed one-by-one and considered as the left-out gene. The remaining genes are used as seed(s) in the RWR algorithms. Depending on the RWR algorithms to be tested, different subsets of these disease and associated genes datasets are extracted as explained on the results sections.

All the network nodes are then scored and ranked according to their proximity to the seed(s). The rank of the disease-gene that was left-out in the current run is recorded. This rank is always between one and the total number of scored genes, minus the number of seeds used for the disease under evaluation. Finally, the Cumulative Distribution Function (CDF) of the ranks of the left-out genes is plotted, as in Mordelet and Vert (2011). It displays the percentage of left-out genes that are ranked within the top k genes. CDFs are used to evaluate and compare the performance of the different algorithms. The plots are focused on the top 60 ranked genes.

#### 2.6.1 Leave-one-out cross-validations on monoplex, aggregated and multiplex networks

For these networks, the seeds used in the RWR algorithms are the gene/protein nodes only. To maximize the size of the test set, we ran the LOOCV with gene-disease associations extracted from DisGeNET v4.0 (Piñero et al., 2016). The DisGeNET dataset contains 6 565 gene-disease associations, corresponding to 4 148 different diseases.

#### 2.6.2 Leave-one-out cross-validation on heterogeneous and multiplex-heterogeneous networks

For these heterogeneous networks, the seeds used in the RWR algorithms are the gene/protein nodes, but also the disease nodes. Given that the nodes in the disease-disease network are OMIM diseases (Hamosh et al., 2005) (material and methods), it is mandatory to use gene-disease associations from OMIM for the LOOCV. The OMIM dataset contains 4 996 gene-disease associations, corresponding to 4 188 different diseases. As in previous applications of the LOOCV, for every disease, each known disease-associated gene is left-out one by one. The remaining disease genes and the disease itself are used as seed nodes. It is to note that in order to simulate an unknown gene-disease association, we also removed the bipartite association linking the left-out gene and the disease of the current run. Doing so, we avoid the artificial top ranking of the left-out genes.

## 3 RESULTS

The main goal of the research presented here was to design a RWR algorithm able to exploit multiple biological interaction sources. We first constructed three biological networks: a protein-protein interaction (PPI) network, a Pathway-derived network and a Co-expression network. We can consider these three networks isolated as monoplex networks. The three monoplex networks can also be merged into an aggregated network. In this case, two proteins A and B can be connected by up to three edges (PPI, Pathways and Co-expression). The aggregated network is composed of 17 559 nodes and 1 659 084 edges (Table 1). In addition, we also considered the 3 networks as layers of a multiplex network. A multiplex network is a collection of networks considered as layers, sharing the same set of nodes, but in which edges belong to different interaction categories.

We also constructed a disease-disease similarity network, in which the nodes correspond to diseases, and the edges to the most significant phenotype similarities between the diseases (materials and methods). Finally, in order to construct a multiplex-heterogeneous network, we linked the disease-disease similarity network to the multiplex network thanks to bipartite gene-disease associations.

We next devised different RWR algorithms, which each leverage the different networks and combinations thereof, and we compared their efficiencies.

### 3.1 Random walk with restart on multiplex networks are more efficient than on monoplex networks

The classical RWR algorithm takes as input a monoplex network. Here, we first adapted the RWR algorithm to navigate a multiplex network (RWR-M). Basically, at each step, the particle can walk from one node to another in the same layer, as in a monoplex network, but it can also move to the same node in another layer of the multiplex network (materials and methods). We compared the performances of the classical RWR and multiplex RWR-M algorithms in retrieving disease-associated genes, thanks to a leave-one-out cross validation (LOOCV) strategy (materials and methods). For that, we created a test set composed of diseases associated to at least two genes in the set of 4 529 protein nodes common to the three networks. This test set contains 273 diseases and 1 312 gene-disease associations. For every disease, each of its associated genes is iteratively left-out, and the remaining gene(s) are considered as seed(s) to run the algorithms. We then compared the ability of the different algorithms to retrieve the left-out gene. Results are displayed in Fig. 2.

**Figure 2:**
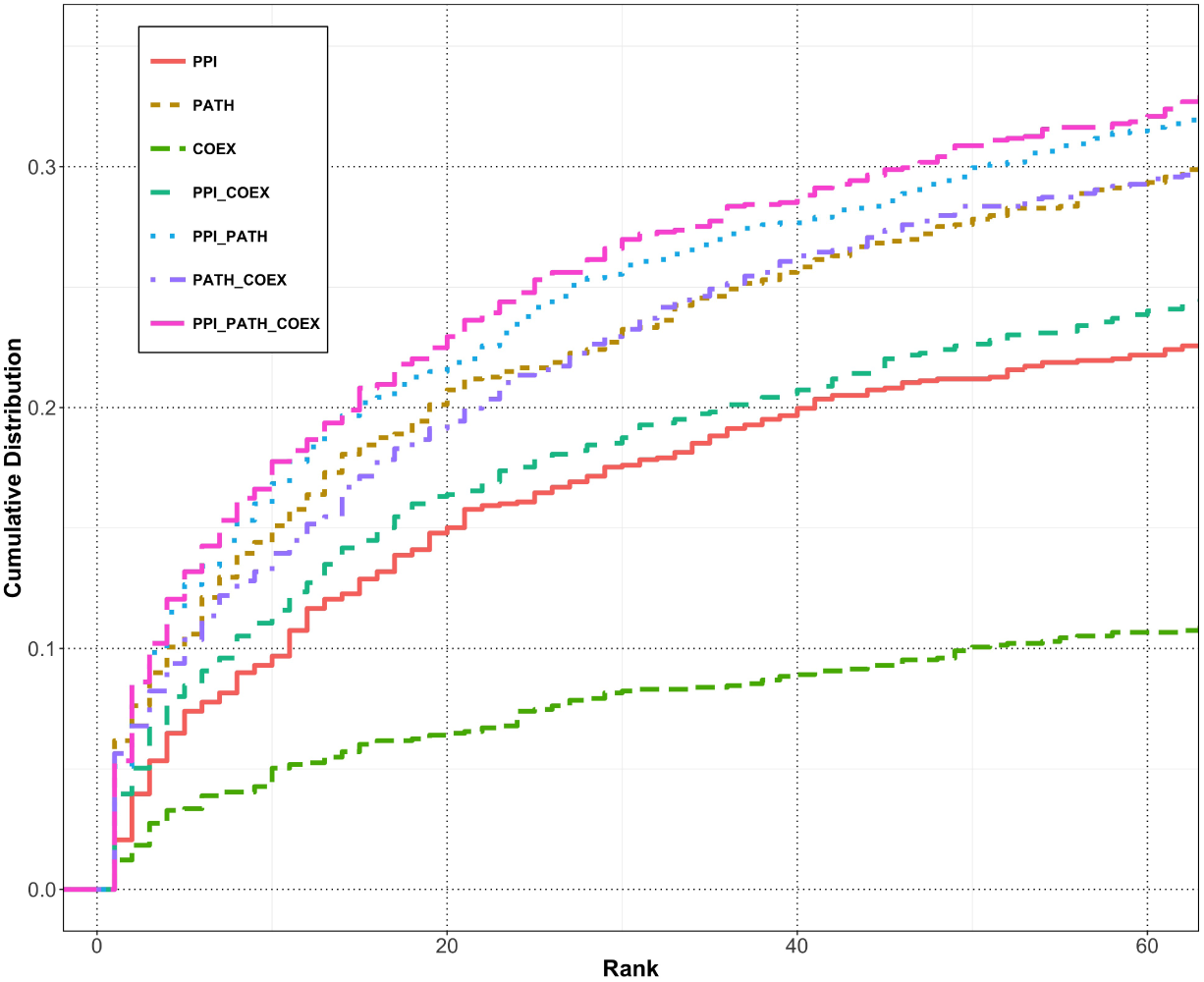
Cumulative distribution functions representing the ranks obtained for the left-out disease genes in the LOOCV with different RWR algorithms. Classical RWR algorithm is applied to the protein-protein (PPI), Pathway (PATH) and Co-expression (COEX) monoplex networks. RWR-M algorithm is applied to combinations of 2 or 3 of these networks, considered as layers of a multiplex network.

Focusing first on monoplex networks, the worst performance is observed for the classical RWR algorithm applied to the Co-expression network. It seems difficult to retrieve known disease-associated genes from a network built from correlations of mRNA expression data alone. The Pathway-derived network achieves the best performance among the monoplex networks, probably because pathways databases are usually built on established biological knowledge and curated.

The RWR-M algorithm, exploiting more than one interaction source in a multiplex framework, performs better than the classical RWR. In particular, despite the low ranking capacities of the co-expression network alone, its integration as a layer in a multiplex framework of two or three layers enhances the performance of the algorithm. Overall, the best ranking result is obtained with the integration of the three network layers (Fig. 2).

### 3.2 Random walk with restart on multiplex networks are more efficient than on aggregated networks

In a second step, we compared the performances of the random walk with restart on multiplex network (RWR-M) with the classical RWR run on the three networks aggregated as a single monoplex network. In the aggregated network, two proteins can be linked by up to three edges (corresponding to the three network sources), and the random walk particle can choose between these different edges to move from its current node to one of its neighbors, as in Li and Li (2012). The ranking ability of RWR-M and classical RWR on the aggregated network are again tested by LOOCV (materials and methods). In this case, we created the test set with diseases associated to at least two nodes in the total of 17 559 nodes corresponding to the union of the nodes of the three networks. The test set contains 537 diseases and 2 892 gene-disease associations.

The ranks of the left-out disease genes with the RWR-M are better than the classical RWR on the aggregated network (Fig. 3). The aggregated and multiplex networks use the same biological data and interaction network sources, but the multiplex framework further keeps tracks of the individual topological structures in each network layer.

**Figure 3:**
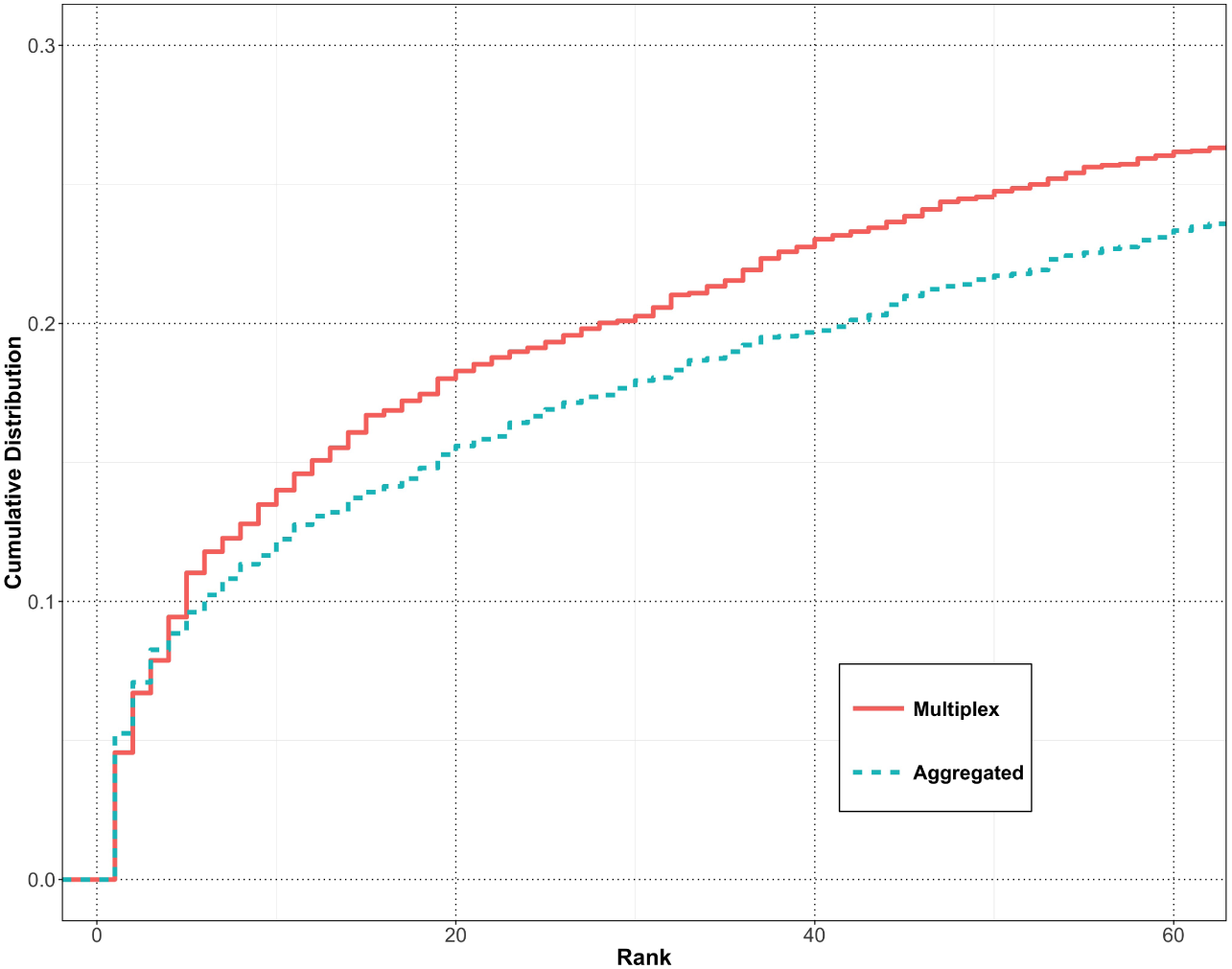
Cumulative distribution functions representing the ranks obtained for the left-out disease genes in the LOOCV with different RWR algorithms. Classical RWR algorithm is applied on the 3 networks aggregated as a single monoplex network, and RWR-M algorithm is applied to combinations of the 3 networks as layers of a multiplex network.

### 3.3 Random walk with restart on multiplex-heterogeneous networks are more efficient than on multiplex or heterogeneous networks alone

We previously compared the performances of RWR algorithms on different combinations of networks containing the same nodes but edges belonging to different interaction categories. The nodes were genes/proteins, and the edges PPI, Pathway and Co-expression interactions. We now wish to extend these comparisons to heterogeneous networks, i.e., networks containing different sets of nodes, such as genes/proteins and diseases.

We first coded the heterogeneous RWR-H algorithm as proposed by Li and Patra (2010) (materials and methods). The RWR-H algorithm takes as input a heterogeneous network composed of a PPI network and a disease-disease similarity network. We constructed the disease-disease similarity network by computing the phenotype similarity between a pair of diseases as the relative information content of their common phenotypes, and linking each disease to its five most similar ones (materials and methods). The PPI and the disease-disease similarity networks are connected by bipartite gene-disease associations. In the RWR-H algorithm, the particle can move from the PPI network to the disease-disease similarity network thanks to these bipartite associations. The conclusion from Li and Patra (2010) was that the RWR-H algorithm on the heterogeneous network performs better than the classical RWR on a monoplex network.

We here compared the ranking capacities of RWR-M and RWR-H by LOOCV. In this case, we created a test set of diseases associated to at least two genes in the set of 12 621 proteins present in the PPI network. The test set contains 242 diseases and 880 gene-disease associations. We can observe first that RWR-M and RWR-H perform better than the classical RWR on the monoplex PPI network (Fig. 4). In addition, the RWR-M algorithm performs slightly better than RWR-H algorithm, since it is able to rank within the top 20 a larger percentage of known gene-disease associations (Fig. 4).

**Figure 4:**
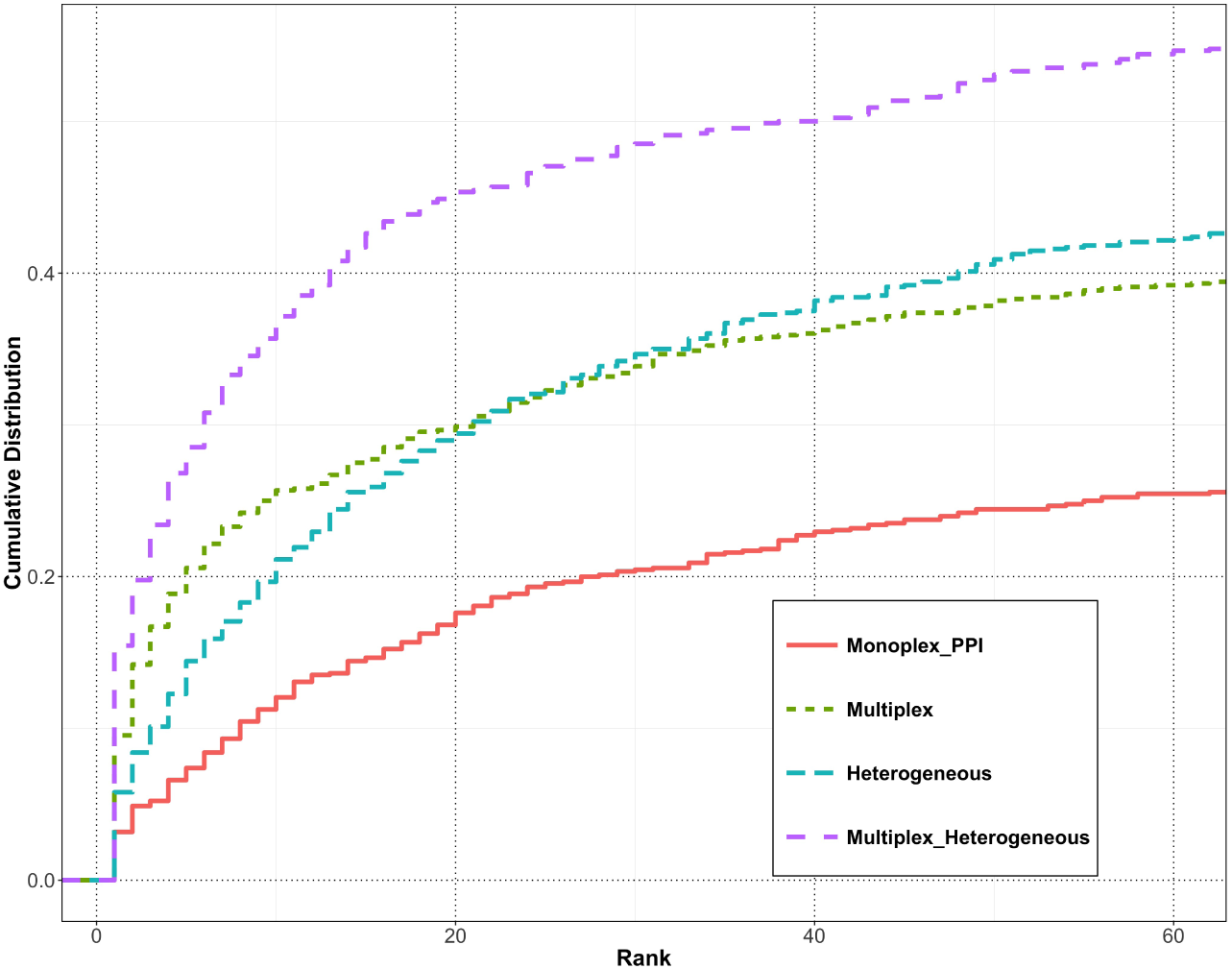
Cumulative distribution functions representing the ranks obtained for the left-out disease genes in the LOOCV with different RWR algorithms. Classical RWR algorithm is applied to the monoplex PPI network, RWR-M is applied to the combinations of the 3 monoplex networks as layers of a multiplex network, RWR-H algorithm is applied to the heterogeneous network composed of the PPI network and the disease-disease similarity network, and RWR-MH algorithm is applied the multiplex-heterogeneous network composed of the 3-layers multiplex network and the disease-disease similarity network.

In this context, an algorithm able to execute a random walk with restart on both multiplex-heterogeneous networks is expected to have better performances. Therefore, we extended our RWR-M approach to heterogeneous networks, defining a random walk with restart on multiplex-heterogeneous networks, RWR-MH (materials and methods). The RWR-MH displays a remarkable amelioration of performances in the prioritization task, since over 45% of the left-out genes are ranked within the top 20 (Fig. 4).

### 3.4 Effect of parameters on the RWR-MH

Finally, we checked the influence of the parameters involved in the RWR-MH algorithm, using again the LOOCV strategy. In this case, we created the test set with diseases associated to at least two genes in the total of 17 559 nodes corresponding to the union of the nodes of the three networks. The test set contains 276 diseases and 1 101 gene-disease associations.

In the applications of the RWR algorithms described previously, the restart parameter was set as *r* = 0.7, as in earlier publications (Li and Patra, 2010; Li and Li, 2012; Zhao et al., 2015; Blatti and Sinha, 2016). Changes in this parameter only slightly affect the results (Fig. 5A).

**Figure 5:**
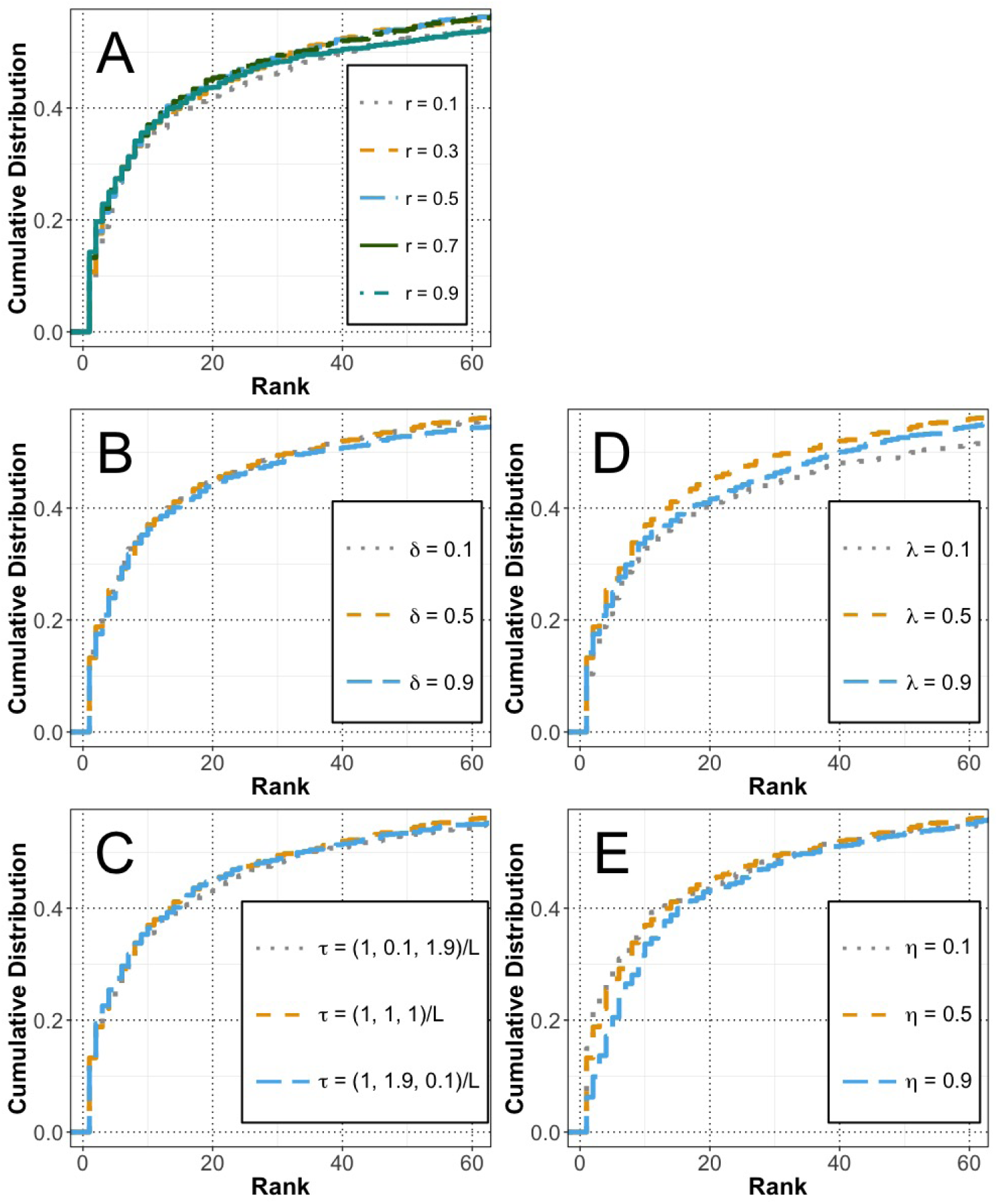
Cumulative distribution function (CDF) of the rank position retrieved for each tested gene by LOOCV when running RWR-MH with variations of the parameters. When one parameter changes, the other parameters remain set on their default value. Variations are tested in: **A)** parameter *r*, **B)** parameter *δ*, **C)** parameter *τ* **D)** parameter *λ* and **E)** parameter *η*.

We then studied the effect of the parameters related to the random walks in multiplex networks, *δ* and *τ*. The parameter *δ* quantifies the probability that the particle jumps from the current node to the same node in a different layer, after a non-restart step. If *δ* = 0 the particle will always stay in the same layer, and if *δ* = 1 the particle will jump to a different layer at each step. However, we did not observe notable changes with variations in this parameter, as displayed in Fig. 5B. The parameter ***τ*** controls the probability of restart in the different layers of the multiplex network. Theoretically, this would allow exploiting our knowledge about the performance of the RWR on the monoplex networks. For instance, it could seem reasonable to favor the restart in the Pathway network and to hinder it in the Co-expression network. However, Fig. 5C does not show notable differences in the performances of the LOOCV with modifications of this parameter.

The parameters used for RWR-H on heterogeneous networks are *λ* and *η*. The parameter *λ* quantifies the probability of jumping between the multiplex and the disease-disease similarity network, using the bipartite gene-disease associations. The larger the value of *λ*, the higher the probability of jumping. If *λ* = 0, the particle does not exploit the bipartite associations between the disease-disease similarity network and the multiplex network. Contrarily, if *λ* = 1, the bipartite gene-disease associations dominate the walks, and the particle is not allowed to deeply explore the topology of each individual network. But variations in this parameter shows only small variations in the performances (Fig. 5D). The parameter *η* quantifies the probability of restart in the multiplex or in the disease-disease similarity network. If *η* = 0, the particle will always restart in the multiplex network. In this case, variations in the parameter slightly influence the performances of the algorithm (Fig. 5E). Overall, the RWR-MH is a very robust algorithm since variations in the parameters do not lead to large variations in the ranking performances.

### 3.5 Examples of application

To illustrate our approach, we applied the RWR-MH algorithm to two different case-examples. We first used the algorithm to predict candidate genes that could be involved in the Wiedemann-Rautenstrauch syndrome, and then explored the network of genes and diseases related to the SHORT syndrome.

#### 3.5.1 Candidate genes for the undiagnosed Wiedemann-Rautenstrauch syndrome

The Wiedemann-Rautenstrauch neonatal progeroid syndrome (WRS; MIM code: 264090) is characterized by intrauterine growth retardation with subsequent failure to thrive and short stature (Toriello, 1990). Patients also display a progeroid appearance, decreased subcutaneous fat, hypotrichosis and macrocephaly (Kiraz et al., 2012). Only a few published cases have been documented, and to our knowledge no gene has been described as causative of the syndrome yet.

To illustrate the application of our approach for disease-associated gene prediction, we applied the RWR-MH algorithm using as seed only the WRS disease node. We then considered the top 25 ranked genes as putative candidates for playing a role in WRS (Fig 6). Many of these top predicted candidate genes, such as *FIG4*, *RNF113A* or *LMNA*, are implicated in diseases directly connected to WRS from phenotype similarities. Mutations in *LMNA* are responsible for the Hutchinson-Gilford Progeria Syndrome (MIM code: 176670) and other premature aging syndromes such as Mandibuloacral Dysplasia with type A Lipodystrophy (MIM code: 248370). However, the targeted sequencing of *LMNA* in few WRS patients did not identify mutations (Kiraz et al., 2012; Hou, 2008). The RWR-MH algorithm also top ranked *ZMPSTE24*, which is known to cause severe progeroid syndromes such as Restrictive Dermopathy (MIM code: 275210) (Navarro et al., 2006). But here also, no mutations were found in 5 WRS patients for this gene (Hou, 2008).

**Figure 6:**
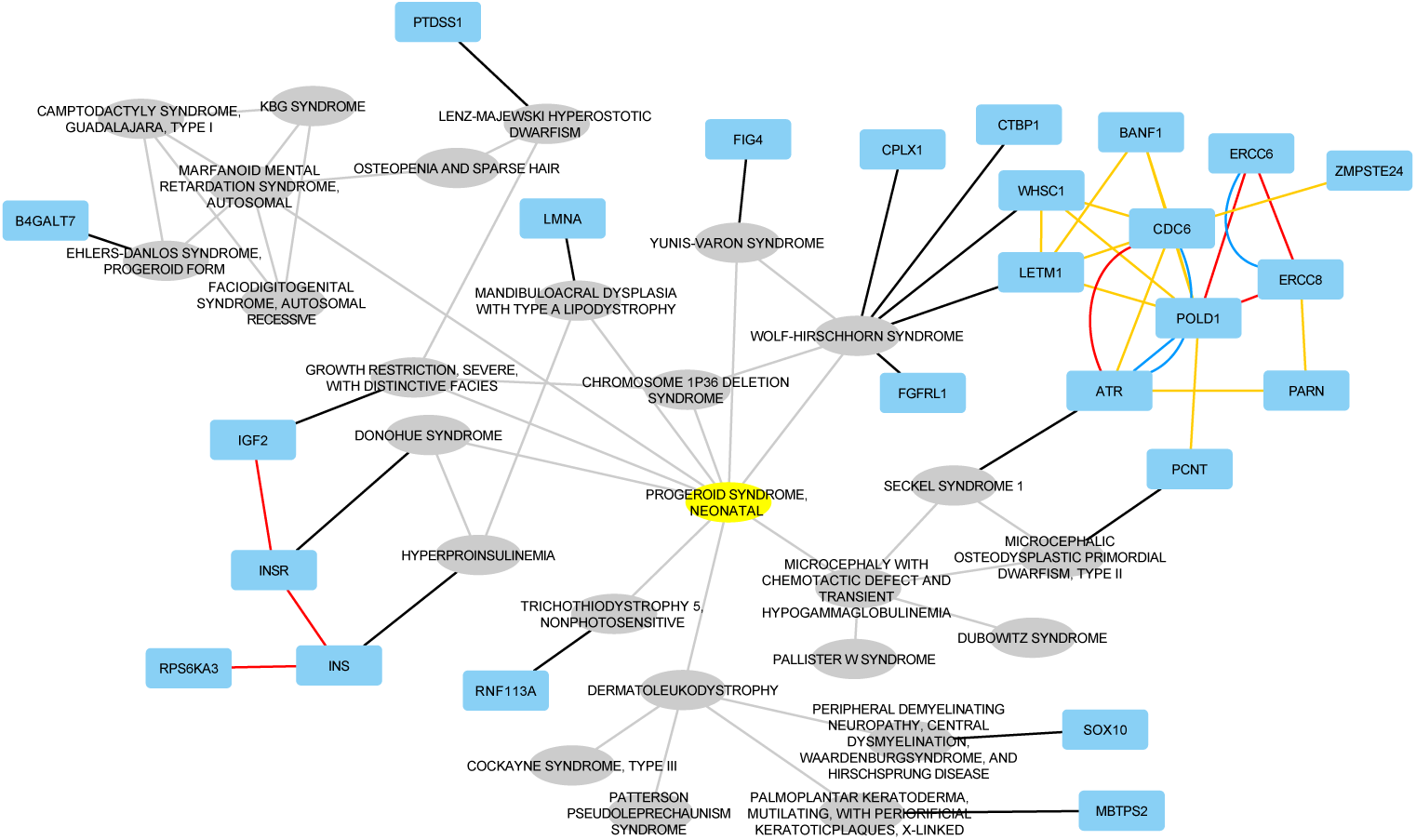
Network representation of the top 25 ranked genes and diseases when the RWR-MH algorithm is executed using WRS as seed (yellow node). Grey elliptical nodes are diseases; Turquoise rectangles are genes/proteins. Black edges are bipartite gene-disease associations from OMIM (Hamosh et al., 2005); Grey edges are the similarity links in the disease-disease network; Blue edges are PPI interactions; Yellow edges are co-expression relationships; Red edges are pathway interactions. The illustration was created using Cytoscape (Shannon et al., 2003). It is to note that results are represented as an aggregated network only for visualization purposes.

Another set of interesting candidates is given by the subnetwork composed of the four genes *IGF2*, *INS*, *INSR* and *RPS6KA3*. All these genes participate in the insulin pathway, and are associated to diseases sharing phenotypes with WRS (i.e., Donohue Syndrome (MIM code: 147670), hyperproinsulinemia (MIM code: 176730), and severe growth restriction (MIM code: 147470)). The insuline pathway is suspected to play a role in WRS (Arboleda et al., 2007). Similarly, a cluster of proteins related to the cell cycle and DNA repair is connected to WRS through the Wolf-Hirschhorn syndrome (MIM code: 194190), and DNA repair defects are also suspected to be involved in WRS (Hou, 2008).

#### 3.5.2 Exploring network vicinity of *PIK3R1* and SHORT Syndrome

SHORT Syndrome (SS; MIM code: 269880) is a rare disease with clinical features defined by its acronym: Short stature, Hyperextensibility of joints and/or inguinal hernia, Ocular depression, Rieger abnormality and Teething delay (Gorlin, 1975). However, these phenotypes do not describe the full range of SS phenotypes, and other clinical features include for instance partial lipodystrophy and insulin resistance (Avila et al., 2016). Mutations in the *PIK3R1* gene are described as the main cause of SS (Dyment et al., 2013; Chudasama et al., 2013; Thauvin-Robinet et al., 2013).

We applied the RWR-MH algorithm using the *PIK3R1* gene and the SS disease as seed nodes, and explored the top 25 ranked diseases and genes, along with their interactions and associations (Fig 7). Many of the top ranked diseases recapitulate phenotypes associated to SS. For instance, permanent neonatal diabetes mellitus (MIM code: 606176) accounts for SS phenotypes associated to insulin resistance. Mandibuloacral dysplasia with type B lipodystrophy (MIM code: 608612) and other diseases associated to lipodystrophy are also top ranked, as well as the growth hormone insensitivity syndrome (MIM code: 262500) that share with SS the phenotypes related to short stature, among others.

**Figure 7:**
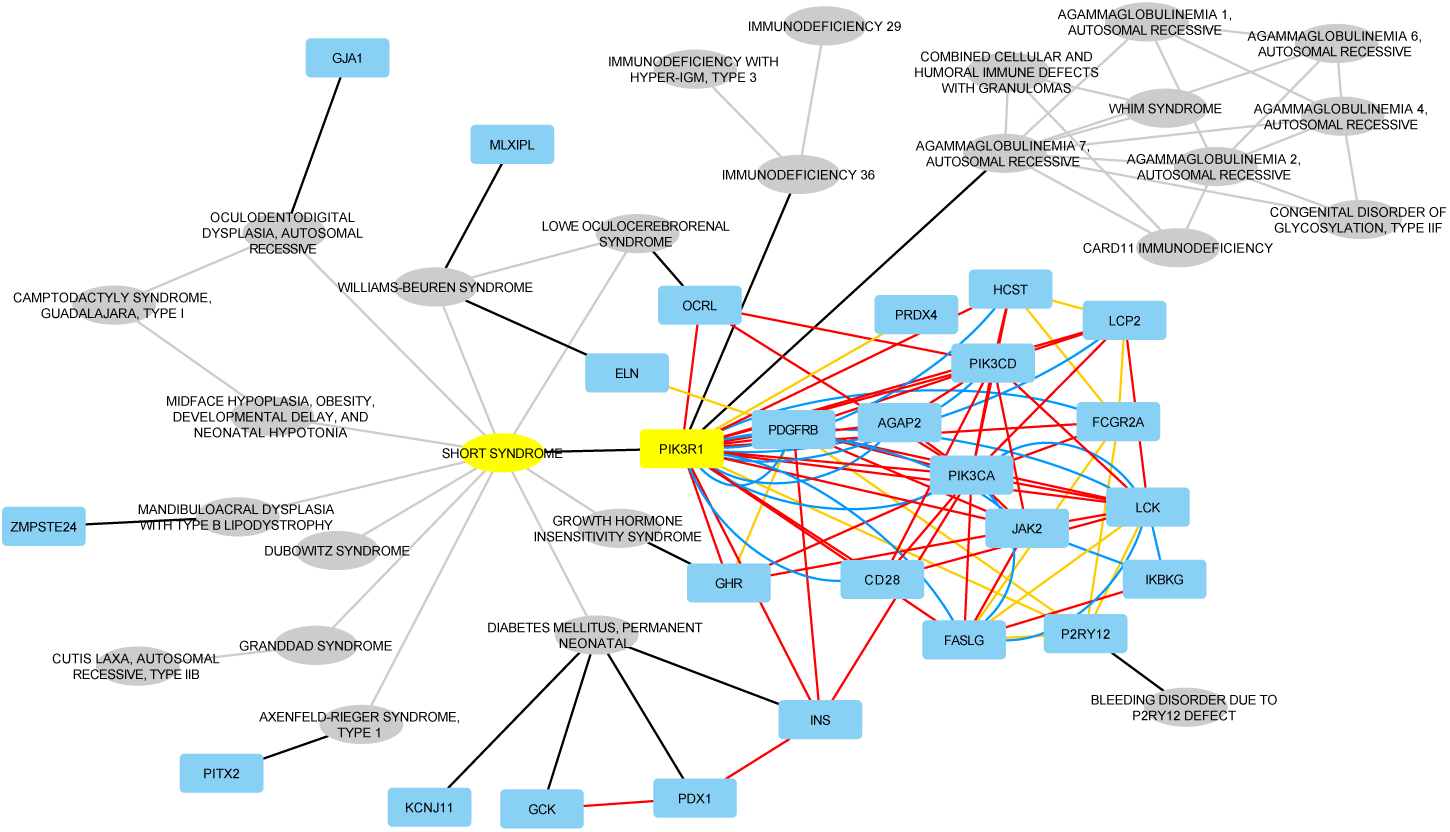
Network representation of the top 25 ranked genes and diseases when RWR-MH is executed using SS as seed disease and *PIK3R1* as seed gene (yellow nodes). Grey elliptical nodes are diseases; Turquoise rectangles are genes/proteins. Black edges are bipartite gene-disease associations from OMIM (Hamosh et al., 2005); Grey edges are the similarity links in the disease-disease network; Blue edges are PPI interactions; Yellow edges are co-expression relationships; Red edges are pathway interactions. The illustration was created using Cytoscape (Shannon et al., 2003). It is to note that results are represented as an aggregated network only for visualization purposes.

Some of the identified subnetworks are very appealing. For instance, we can observe a loop linking the SS, its associated gene, *PIK3R1*, the Lowe oculocerebrorenal syndrome (MIM code: 309000) and its associated gene *OCRL*. These two diseases share a noticeable amount of phenotypes, including growth retardation and glucose intolerance. The *PIK3R1* and *OCRL* genes are coding proteins involved in the same pathway: synthesis of phosphatidylinositol phosphates at the plasma membrane (Reactome code: R-HSA-1660499). Therefore, we can hypothesize a common deregulation of this pathway in the two diseases, leading to shared phenotypes.

Similarly, we can point to the subnetwork containing the *ELN* gene, implicated in the Williams-Beuren Syndrome (MIM code: 194050). Many phenotypes associated to this syndrome are similar to SS and Lowe oculocerebrorenal syndrome. In this case, the *ELN* gene is linked to the *PDGFRB* gene by a co-expression relationship. *PDGFRB* is highly connected to many nodes in the subnetwork, including to *PIK3R1*, by pathway interactions. The co-expression interaction between *PDGFRB* and *ELN* is intriguing because the two genes are, to our knowledge, not described to be involved in the same pathway or process. However, they seem to be regulated by the same microRNA-29 family (Zhang et al., 2012; Cushing et al., 2015).

Overall, these results could also allow pointing to other candidate genes, predicted to be involved in the SS. This is interesting as, for instance, Dyment et al. (2013) did not find any mutation in the *PIK3R1* gene in one of the seven tested patients.

## 4 DISCUSSION

Physical and functional relationships between genes and proteins are diverse. They are identified or derived from various approaches, each having its own features, strengths and weaknesses. In this context, the integration of different sources of interaction, exploiting data pluralism, is expected to outperform current approaches dealing with single networks. Indeed, the combination of different large-scale interaction datasets increases the available biological information, and potentially reduce the bias and incompleteness of isolated sources (Menche et al., 2015).

We and others also hypothesized that the multiplex framework, which retains information on the topology of the individual networks, would perform better as compared to the aggregation of the different interaction sources, (Kurant and Thiran, 2006; Kivelä et al., 2014; Battiston et al., 2014; Didier et al., 2015). We have shown previously, for instance, that the multiplex framework is more efficient than network aggregations to extract communities from biological networks (Didier et al., 2015). We extended here the RWR algorithm by designing the RWR-M algorithm able to leverage multiplex networks. The performances of the RWR-M algorithm are clearly improved as compared to previous algorithms navigating monoplex networks, such as RWR on PPI networks (Köhler et al., 2008), or RWR on aggregated networks (Li and Li, 2012). It is particularly interesting to note that even if a monoplex network, such as the co-expression network, displays poor ranking performances isolated, its integration as a layer of a multiplex network leads to an increase of the performance, thereby demonstrating the potential of the RWR-M strategy.

Moreover, we extended our algorithm to integrate multiplex-heterogeneous networks. To this goal, we first built a disease-disease similarity network based on the information content (IC) of the shared phenotypes between every pair of diseases. Previous approaches building disease-disease networks, such as the ones proposed by Li and Patra (2010); Li and Li (2012), were based on MimMiner (van Driel et al., 2006). MimMiner mines OMIM full-text and clinical synopsis to compute similarity between diseases. Contrarily, our approach is based on the controlled classification of phenotypes in an ontology, and considers both the ontological structure and the frequencies of phenotypes.

We evaluated the algorithms with a Leave-One-Out Cross Validation (LOOCV) strategy, using a cumulative distribution function (CDF) to display the results. As compared to a more classical Receiver Operating Curve (AUC), as detailed for instance in Mordelet and Vert (2011), the CDF ranks all the nodes in the networks. In our case, this means that the total of 17 559 nodes are ranked, even if the plots focus on the top 60. Contrarily, previous approaches were ranking a subset of genes related to the left-out gene, for instance the top 100 closest genes in the genome (Köhler et al., 2008; Li and Patra, 2010; Li and Li, 2012; Zhao et al., 2015). CDF thereby results in a more general validation than other methods.

Thanks to the LOOCV, we demonstrated that when the RWR algorithm is applied on this complex multiplex-heterogeneous network, an approach that we called RWR-MH, the prioritization results are far better than those of all other versions of the algorithm. We have also demonstrated that the RWR-MH algorithm displays a robust behavior upon variations of the different parameters, which are globally inducing no or only few changes in the results. This was previously observed for variations in the parameters of a RWR-H algorithm (Li and Patra, 2010; Zhao et al., 2015). However, it is to note that, although the global curves of the LOOCV CDF do not change significantly when parameters vary, a focused analysis and network representation of the top 25 ranked genes and diseases in a real-case applications would reveal variations. In these applied cases, changes in parameters can be used to tune the output. For instance, the parameter ***τ*** would allow giving more emphasis on some input network layers, based on prior knowledge related to their biological relevance.

Random walks with restart in biology have been applied to predict disease-associated genes (Köhler et al., 2008; Li and Patra, 2010; Li and Li, 2012; Zhao et al., 2015; Xie et al., 2015), but also to predict drug-target interactions (Chen et al., 2012; Liu et al., 2016) and adverse drug reactions (Chen et al., 2016), and to identify clusters from PPI Networks (Macropol et al., 2009). Smedley et al. (2014, 2015) developed Exomiser, where RWR is applied to prioritize genes and variants in the context of whole-exome sequencing. We applied here our advanced version of the random walk with restart algorithm, RWR-MH, to two real-case biological examples. In the first one, we predicted candidate genes that could be associated to the WSR syndrome, whose responsible gene(s) remain to be described. We hereby demonstrate the usefulness of the approach to study disease etiology and help diagnose patients. The next step will be to validate these predictions, for instance using exome-sequencing data. We also applied the RWR-MH algorithm to study the network vicinity of a disease, the SHORT syndrome, and its associated gene, *PIK3R1*. We show that the disease is sharing phenotype with other syndromes, which are caused by genes in the neighborhood of *PIK3R1* when multiple interaction types are considered. This is an additional example of the fact that mutations in genes participating to the same pathway, or more generally biological processes, lead to diseases with similar phenotypes (Oti et al., 2006).

The main underlying hypothesis of the work presented here is that the integration of multiple interaction sources, each having its own features and biases, will improve the results of the random walks by providing complementary data. For instance, in the application of the RWR-MH to the WRS syndrome, we retrieved as top candidates the *LMNA* and *ZMPSTE24* genes. The *ZMPSTE24* gene codes a peptidase acting during the post-translation modifications of the prelamin A, coded by *LMNA*, to undergo the complete maturation to lamin A. It is interesting to note that the direct interaction between the products of *LMNA* and *ZMPSTE24* is missing in the databases we used to construct the multiplex network. However, the *ZMPSTE24* node is retrieved through different trajectories in the random walk. Hence, the combination of multiple network sources in this case allow completing missing interaction data.

We focused our applications on a multiplex network composed of a PPI, a Pathway and a Co-Expression network. Other biological networks could be collected or constructed from -omics data, and integrated into our multiplex-heterogeneous framework. Functional interactions can be derived, for instance, by connecting genes annotated for the same Gene Ontology (GO) terms (Ashburner et al., 2000). It would also be valuable to include networks with transcription factors - targets genes, non-coding RNAs, as well as drug and therapeutic targets.

The highly connected nodes, called hubs, can be genes or proteins highly connected and central in the cells, but can also result from biased biological experiments studying “fashion” proteins, such as *TP53* in cancer or *APP* in Alzheimer. RWR algorithms and other network propagation algorithms are biased towards highly connected proteins, as demonstrated by Erten et al. (2011). In this context, poorly-connected and unwell-known nodes, which are also potentially relevant for diseases, are more complicated to find than highly-connected and well-known proteins. To address this issue, biased random walks have been developed to favor the walk of the particle according to network topological features (Battiston et al., 2016). In the simplest case, the transition probability depends on the degree of the neighbors of the current node: the walk of the particle can be tuned towards less connected nodes (Bonaventura et al., 2014). Such a degree-biased random walk could be applied to the RWR-MH algorithm in the future.

In addition, for the sake of simplicity, all the networks considered in this study are unweighted. Nevertheless, the extension to weighted networks is straightforward, as pointed out in the material and methods. The use of weighted networks could improve the prioritization results because we can assign larger transition probabilities to the most confident interactions or to the more similar diseases. For instance, STRING database stores scored protein-protein interactions indicating its confidence based on the evidences (Szklarczyk et al., 2015). The edges in our Co-expression network are established based on threshold imposed on the value of the computed correlation coefficient. This coefficient can be included into the Co-expression network to favor the transitions between the proteins whose expressions are more correlated. In addition, we built the disease-disease similarity network according to the similarity scores between every pair of diseases. This score could be introduced into the corresponding edges.

## 5 ACKNOWLEDGEMENTS

## 7 AUTHORS CONTRIBUTIONS

